# Coculturing bacteria leads to reduced phenotypic heterogeneities

**DOI:** 10.1101/423715

**Authors:** Jasmine Heyse, Benjamin Buysschaert, Ruben Props, Peter Rubbens, Andre G. Skirtach, Willem Waegeman, Nico Boon

## Abstract

Isogenic bacterial populations are known to exhibit phenotypic heterogeneity at the single cell level. Because of difficulties in assessing the phenotypic heterogeneity of a single taxon in a mixed community, the importance of this deeper level of organisation remains relatively unknown for natural communities. In this study, we have used membrane-based microcosms that allow the probing of the phenotypic heterogeneity of a single taxon while interacting with a synthetic or natural community. Individual taxa were studied under axenic conditions, as members of a coculture with physical separation, and as a mixed culture. Phenotypic heterogeneity was assessed through both flow cytometry and Raman spectroscopy. Using this setup, we investigated the effect of microbial interactions on the individual phenotypic heterogeneities of two interacting drinking water isolates. We have demonstrated that interactions between these bacteria lead to an adjustment of their individual phenotypic diversities, and that this adjustment is conditional on the bacterial taxon.

**Importance:** Laboratory studies have shown the impact of phenotypic heterogeneity on the survival and functionality of isogenic populations. As phenotypic heterogeneity is known to play an important role in pathogenicity and virulence, antibiotics resistance, biotechnological applications and ecosystem properties, it is crucial to understand its influencing factors. An unanswered question is whether bacteria in mixed communities influence the phenotypic heterogeneity of their community partners. We found that coculturing bacteria leads to a reduction in their individual phenotypic heterogeneities, which led us to the hypothesis that the individual phenotypic diversity of a taxon is dependent on the community composition.

## Introduction

Genetically identical bacteria are known to exhibit single cell heterogeneity under controlled laboratory conditions (1–3). These heterogeneous traits include morphological traits, such as cell size, as well as biochemical properties, such as protein and mRNA content. The individualisation of identical sister cells in clonal populations occurs rapidly after cell division (4). Cells can be partitioned into clusters of cells with similar traits, called phenotypes. The variation in phenotypes within sympatric isogenic populations is referred to as the phenotypic heterogeneity (5).

Noise in gene expression is known to be one of the main drivers of phenotypic heterogeneity (6–8). At first glance, a heterogeneous gene expression appears to be disadvantageous, as it may reduce the mean fitness of the population under the prevailing environmental conditions (9). However, several studies have indicated that biological noise is an evolved and regulated trait (10, 11), which offers benefits for the survival (12, 13) and functionality (14–16) of a clonal population. The aforementioned studies have revealed that isogenic bacterial populations are not homogeneous populations. Instead, they behave as communities consisting of phenotypic subgroups, which may differ in quantitative (*i.e.* continuous variation in phenotypic traits) and qualitative (*i.e.* distinct phenotypic states) aspects.

In nature, bacteria are not encountered as isolated populations, but they are a part of a larger association where many microorganisms coexist. To date few research has been devoted to the occurrence and functional consequences of phenotypic heterogeneity in natural, mixed communities (17, 18) and our knowledge regarding factors that influence phenotypic heterogeneity is limited. One of the reasons for this is that it is difficult to assess the heterogeneity of a single taxon within a mixed community. Recently, several experimental approaches that assess the metabolic diversity of a single taxon in natural communities have been developed (19, 20). However, these approaches rely on FISH-probes that bind to 16S rRNA gene sequences for identification of the taxon of interest. Hence, they do not allow to exclude the possibility that some of the observed phenotypic differences are caused by minor genetic differences between bacteria with very similar 16S rRNA genes.

Two laser-based methods that are suitable for assessing phenotypes are flow cytometry and Raman spectroscopy (21–23). Two types of light can be detected by the flow cytometer, that is scattered light and fluorescence. The scattered light provides information about the basic characteristics of the cells (*e.g.* size, shape and surface properties), while the fluorescence data provides additional information about the cell properties for which it has been stained (*e.g.* nucleic acid content, metabolic activity, etc.) (24). Flow cytometry thus gives information regarding morphological as well as specific physiological properties of single-cells. The Raman spectrum of a single cell consists of a combination of the individual spectra of all the compounds that make up this cell (*e.g.* proteins, nucleic acids, fatty acids, etc.). This results in a complex spectrum, which can be interpreted as a chemical fingerprint of the cell (25, 26). Hence, single-cell Raman spectra offer an in depth view on the biochemical composition of each phenotype.

A tool that can help to answer questions that are difficult to study directly in natural communities is a synthetic ecosystem. A synthetic ecosystem consists of a selected set of species under specific conditions. They are controllable and have a reduced complexity in comparison to natural communities (27). Hence, they provide a way to test ecological theories in order to better understand the rules of nature (28). A specific setup for these synthetic ecosystems are co-cultures. The principle of such a system is that two or more bacterial populations are cultivated together with some degree of contact between them, which allows to study their interactions (29).

An unanswered question, and the focus of this study, is whether bacteria in mixed communities influence the phenotypic heterogeneity of their community partners. Here, we used a synthetic community setup where two isolates were used as model organisms. Four synthetic communities were created. The isolates were grown in axenic cultures as a reference for non-interacting genotypes. To be able to study the individual community members separately after they have been interacting via their joint medium, a coculture with physical separation by a membrane was created. Lastly, a mixed culture without physical separation, representing ‘full interaction’, was created. Phenotypes were assessed through flow cytometry and single-cell Raman spectroscopy. Furthermore, we applied and evaluated a novel machine learning approach to quantify synthetic community composition through flow cytometric fingerprinting.

## Results

We aimed to evaluate whether the phenotype and phenotypic heterogeneity of a single taxon in a dual-species coculture is mediated by interactions with a partner taxon. Two drinking water isolates, an *Enterobacter* sp. and a *Pseudomonas* sp., were used as model organisms. The experimental design consisted of four synthetic communities: two axenic cultures, a coculture with physical separation between the taxa (partial interaction), and a mixed culture (full interaction) (**Fig. 1**). The synthetic communities were monitored for 72 h. Every 24 h population phenotypic diversity was assessed by flow cytometry. At 72 h, populations were analysed using single-cell Raman spectroscopy. Cell viability throughout the experiment was verified through SGPI staining (**Fig. S4**). Cell populations remained viable throughout the course of the experiment and viability was found to be similar between the cocultures and axenic cultures (**Fig. S4**). In the following results the physically separated culture is referred to as the ‘coculture’, while ‘mixed culture’ indicates the culture without physical separation.

**FIG 1.**
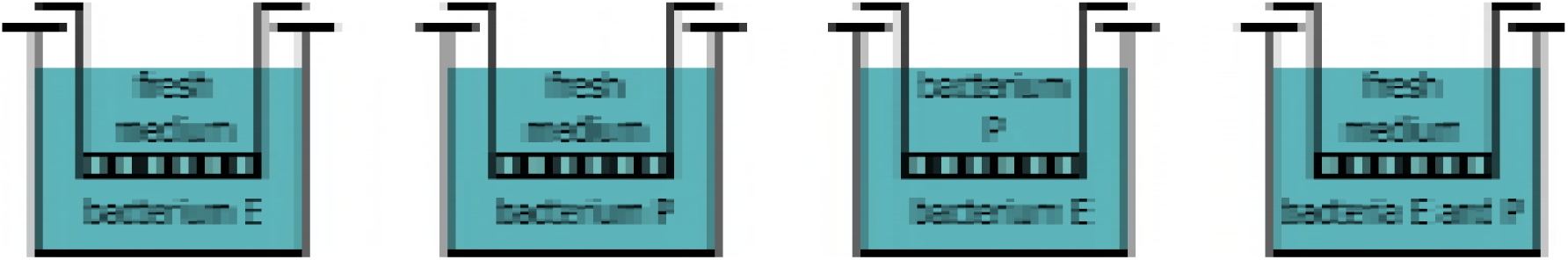
Illustration of the experimental setup. Bacteria in apical and basal phase can interact via metabolites in their shared medium while they are physically separated by the membrane of the cell culture inserts. Four synthetic communities were created: two axenic cultures, a coculture and a mixed culture. There were biological replicates (n = 3) for each synthetic community.

### Flow cytometric diversity assessment

To evaluate whether microbial interactions can lead to changes in the phenotypic heterogeneity of interacting organisms, cytometric diversity estimates were used as measures of phenotypic heterogeneity. For this, an equal spaced binning grid was used to arbitrarily split up the cytometric parameter space in operational phenotypic units. The signals of both scatter and fluorescence detectors were used, implying that the diversity is a measure of population heterogeneity in terms of both morphological traits and nucleic acid content. Note that the calculated diversity metrics are independent of the taxon abundances (**Fig. S5**), as all populations were subsampled to equal cell counts prior to diversity estimation.

The phenotypic community structure was first investigated through an alpha-diversity (*i.e.* within sample diversity) assessment. For both taxa, the diversity of the individual taxon was larger when present in the axenic culture compared to when the same taxon was present as a member of the coculture. Not only the phenotypic diversity (D_1_ and D_2_,), which include both richness and evenness, decreased (**Fig. S6**), but the phenotypic richness (D_0_) of the coculture members decreased as compared to the axenic cultures (**Fig. 2A**). This indicates that the interaction did not only lead to a reorganization of the phenotypic community structure (*i.e.* change in the relative abundances of the cytometric bins), but that the number of non-empty bins on the cytometric fingerprint was reduced due to the interaction, implying not only a redistribution of trait abundance, but a reduction in trait heterogeneity.

**FIG 2.**
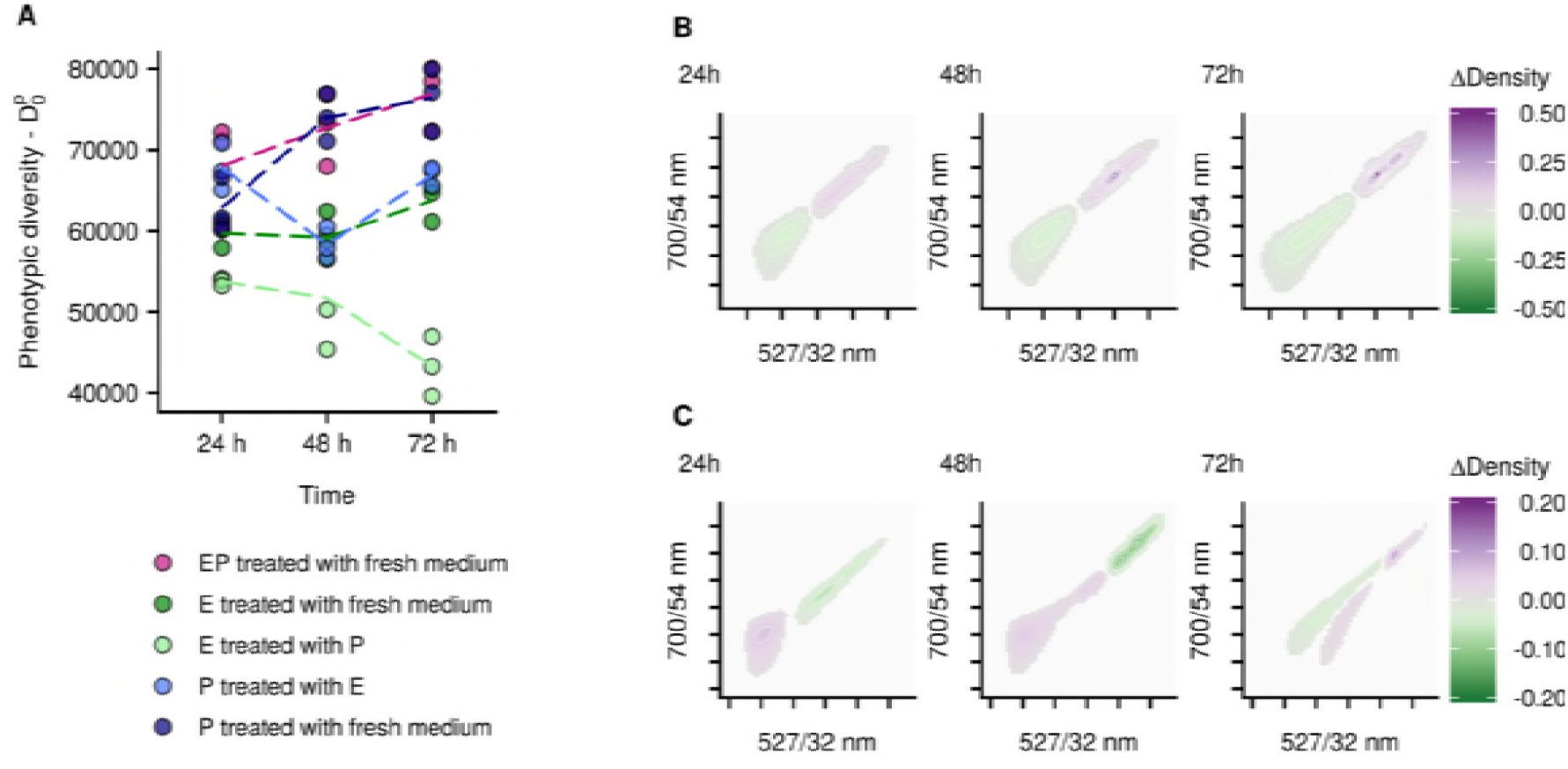
Phenotypic alpha diversity D_0_ for both individual bacterial taxa in communities of axenic cultures, cocultures and mixed cultures **(A)**. The taxa are denoted as taxon E *(Enterobacter* sp.) and P (*Pseudomonas* sp.), respectively. The populations are indicated with names in the form of ‘X treated with Y’, where X is the taxon in the sample (E, P or EP) and Y is what was present on the other side of the membrane (E, P or fresh medium). There were biological replicates (n = 3) for each community. The dashed lines indicate the average trend of the replicates. Contrast analysis of the phenotypic fingerprints to compare the phenotypic community structure of axenic cultures and coculture members with respect to fluorescence intensity. Each plot is a comparison between the axenic culture and coculture of the same taxon at the same time point, averaged over the three biological replicates. The colour gradient indicates whether density in the coculture increased (purple) or decreased (dark green) relative to their respective axenic culture at the specified time point. The upper row presents contrast results for *Enterobacter* **(B)**. The lower row presents contrast results for *Pseudomonas* **(C)**. If the difference between the two communities is lower than 0.01 no contrast value is shown on the graphs, which causes the appearance of different cluster. Note that different scales were used for the different taxa.

Using a contrast analysis, differences between the phenotypic fingerprints of populations can easily be visualised in bivariate parameter spaces. To evaluate whether the observed lower diversities were linked with specific shifts in the cytometric fingerprint, differences in scatter and fluorescence patterns of the axenic cultures and the cocultures were assessed. The differences in scatter patterns were limited for both taxa (**Fig. S10**). In contrast, a clear difference in fluorescence intensity was observed (**Fig. 2, B and C, Fig. S7**). For *Enterobacter* there was a shift towards high fluorescence cells in the coculture as compared to the axenic culture. This difference became larger over time. For *Pseudomonas* there was a more limited difference, with a small enrichment of lower fluorescence cells. Thus, there was not only a reduction in population diversity, but there was also a shift of the population fingerprint. Moreover, this shift was taxon-dependent.

To further compare the cytometric fingerprints of the different populations, a PCoA ordination was generated based on the Bray-Curtis dissimilarities between the fingerprints (**Fig. 3**). In this ordination, the fingerprints of the taxa, both under axenic and under coculture growth, are separated, with the mixed culture in between. The populations show a significant shift in their phenotypic structure through time (p = 0.001, r^2^ = 0.154). In addition, there is a significant difference in the fingerprint of *Enterobacter* when present as an axenic culture compared to being present in the coculture (p = 0.001, r^2^ = 0.455). For *Pseudomonas* the differences between the axenic cultures and coculture members were not significant (p = 0.092, r^2^ = 0.170). The mixed culture shifted from a community that is more resembling *Enterobacter* at the first measurement, towards a community that is more similar to *Pseudomonas* at the second and third measurement.

**FIG 3.**
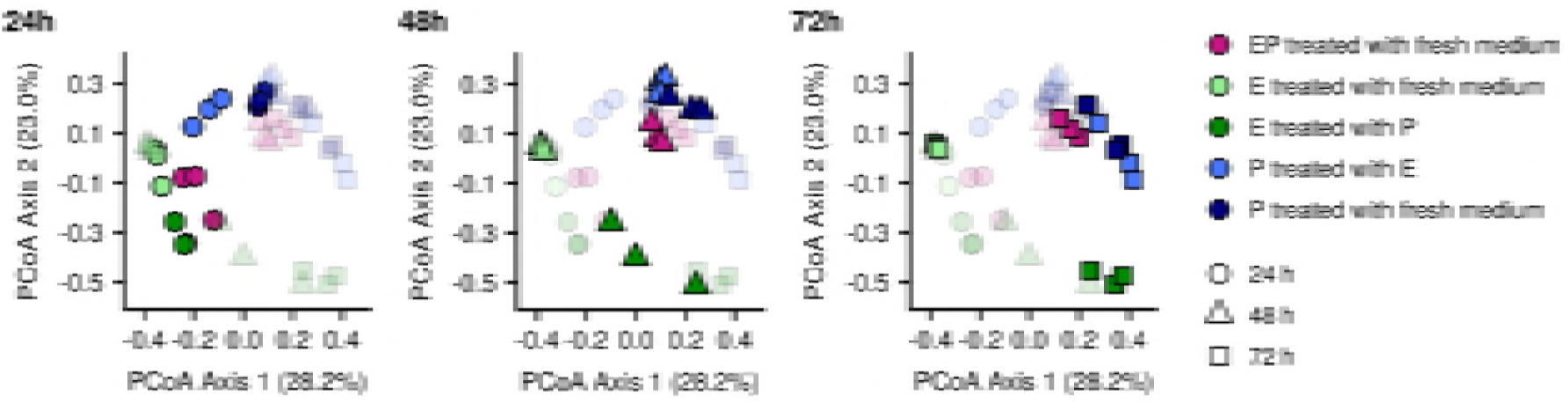
PCoA ordination of the Bray-Curtis dissimilarities between the phenotypic fingerprints for both individual bacterial taxa in communities of axenic cultures, cocultures and mixed cultures. The ordination is shown in three graphs, split according to time, since this allows for easier interpretation of how the different communities are relating to each other at each time point. The taxa are denoted as taxon E (*Enterobacter* sp.) and P (*Pseudomonas* sp.), respectively. The populations are indicated with names in the form of ‘X treated with Y’, where X is the taxon in the sample (E, P or EP) and Y is what was present on the other side of the membrane (E, P or fresh medium). There were biological replicates (n = 3) for each community.

To better understand the interaction that was occurring between *Enterobacter* and *Pseudomonas*, we applied a novel machine learning approach to infer the relative abundances of both taxa in the mixed community. Previous results confirmed our initial hypothesis that the phenotypic diversity of a taxon can be influenced by the presence of other taxa. In order to take this into account, a random forest classifier was trained, for each time point separately, on the fingerprints of the coculture members at the corresponding time point, as these are expected to be the most biologically accurate (Supplementary Results and Discussion). The predictions indicate a higher abundance of *Enterobacter* in the community at 24 h, followed by a gradual enrichment of *Pseudomonas* at the second and third time point (**Fig. 4**).

**FIG 4.**
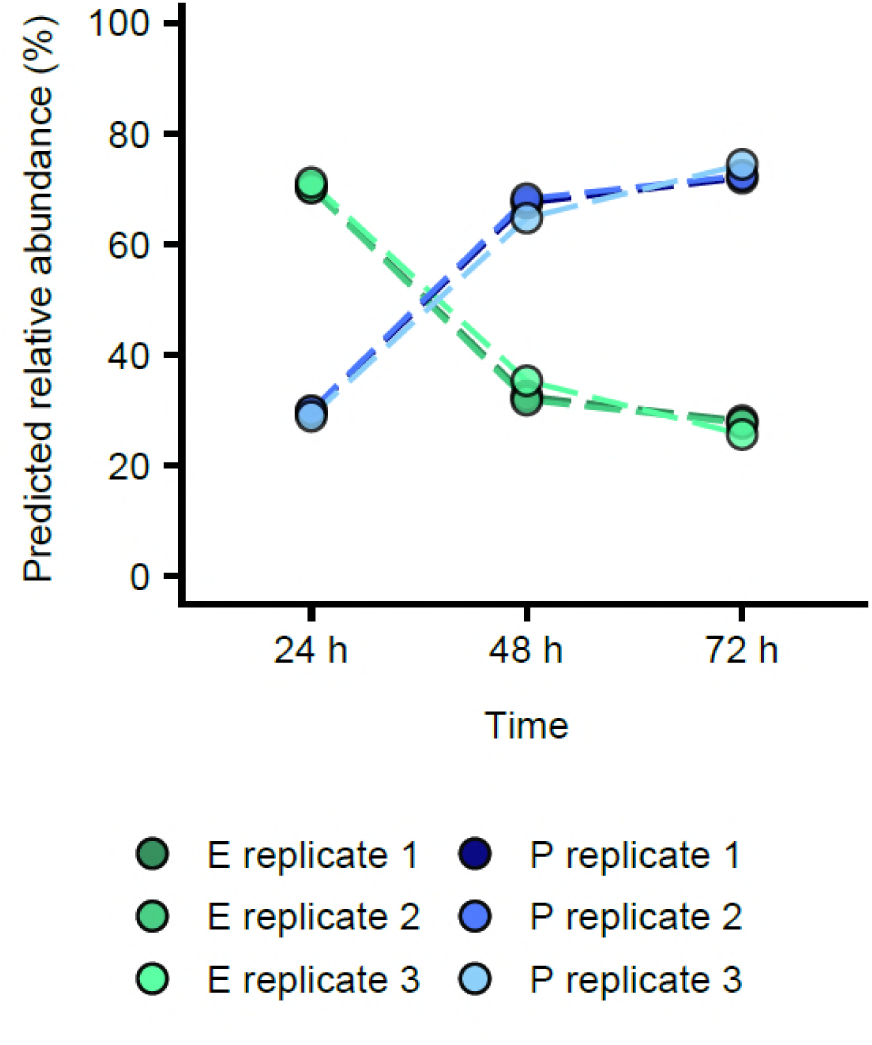
Predicted relative abundances in the mixed cultures. The random forest classifiers that were used to infer community composition were constructed using the fingerprints of the coculture members at the corresponding time point as input data. Green lines indicate the predicted relative abundances of *Enterobacter*, blue lines indicate the predicted relative abundances of *Pseudomonas*. The different shades correspond to biological replicates (n = 3).

In summary, both *Enterobacter* and *Pseudomonas* showed lower phenotypic diversities in the coculture compared to their axenic culture counterparts. However, while the overall phenotypic community structure did not change substantially for *Pseudomonas* (*i.e.* small differences in beta-diversity and limited shift towards lower fluorescence intensity cells), there was a clear shift in the phenotypes of the *Enterobacter* population (*i.e.* large differences in beta-diversity and a clear shift towards higher fluorescence intensity cells).

### Raman phenotyping

The cytometric phenotype only takes into account the morphological characteristics and nucleic acid content of the cells. However, phenotypes can differ in more cell constituents than nucleic acids alone. The Raman spectrum of a single cell offers a more in depth view on the biochemical phenotype compared to flow cytometry. Raman spectroscopy was used to measure single cell spectra for each of the populations of *Enterobacter* and *Pseudomonas* in the axenic cultures and the coculture at 72 h.

The spectra hold 333 wavenumbers over the selected biologically relevant range. To gain insight in the separability of cells from the different populations, spectra were visualised through PCA after preprocessing of the data (see materials & methods) (**Fig. 5 A**). The spectra of the *Enterobacter* populations were clearly separated. A large overlap between the spectra of *Pseudomonas* that was grown in axenic culture and *Pseudomonas* that was grown in the coculture was observed. However, when performing PCA for each taxon separately, cells from each synthetic community were separated well (**Fig. 5 B and C**). This confirms the previous results, indicating that for both taxa a phenotypic shift occurred, but that this shift was larger for *Enterobacter* than for *Pseudomonas*.

**FIG 5.**
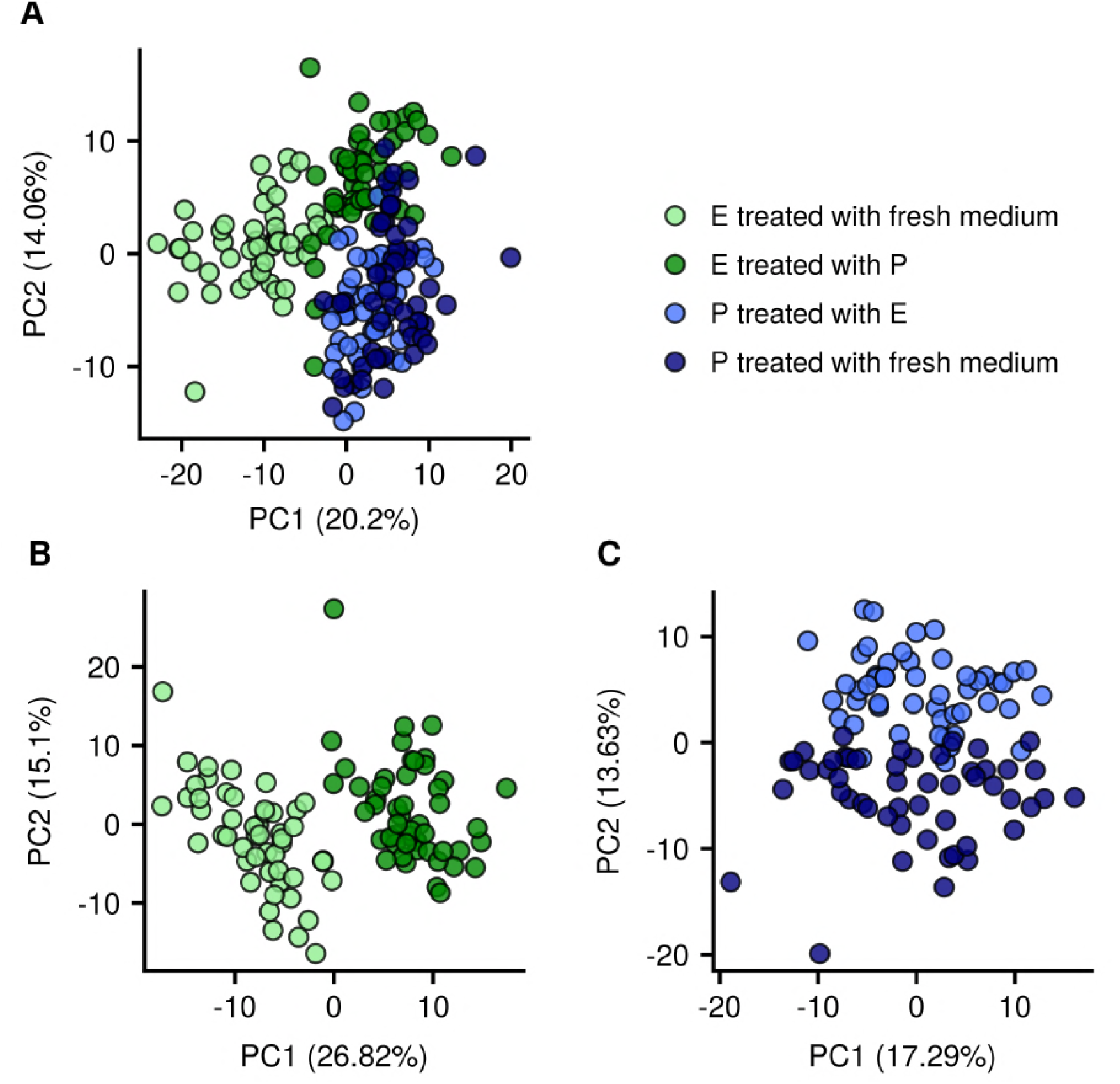
Visualisation of the separability of the single cell Raman spectra for *Enterobacter* and *Pseudomonas* in axenic culture and coculture at 72 h. There are 51 single cell measurements for each population. The taxa are denoted as taxon E (*Enterobacter* sp.) and P (*Pseudomonas* sp.), respectively. The populations are indicated with names in the form of ‘X treated with Y’, where X is the taxon in the sample (E, P or EP) and Y is what was present on the other side of the membrane (E, P or fresh medium). PCA was carried out for the entire dataset **(A)**, for the spectra of *Enterobacter* separately **(B)** and for the spectra of *Pseudomonas* separately **(C)**.

Since the Raman spectrum of a single cell is a combination of the spectra of all compounds that make up this cell (*e.g.* proteins, nucleic acids, fatty acids, etc.), the signal intensity at every wavenumber is the result of all compounds that produce a signal at this wavenumber. The Raman spectra of all DNA and RNA bases are available from literature (42) as well as information regarding peak regions that are assumed to be related to nucleic acids in general (43). We aimed to investigate whether the shift in fluorescence intensity that was observed through flow cytometry was caused by a changing DNA or RNA content, and in this way get more information about the cause of the observed phenotypic shift. Based on this tentative peak assignment, differences in nucleic acids between the coculture and the axenic populations were observed for both taxa (**Fig. S8**). However, there was no consistency in whether this considered an increase or a decrease (*i.e.* for some wavenumbers the average intensity was higher in the coculture, while for other wavenumbers the intensity was higher in the axenic culture). When considering only uracil and thymine it remained impossible to draw a conclusion regarding whether DNA or RNA differences contributed most to the observed phenotypic shift (**Fig. S8**).

## Discussion

There is an interest in understanding the implications of phenotypic heterogeneity in both natural and engineered microbial ecosystems. Our current knowledge is mainly based on experimental set-ups using axenic cultures. This is partly due to the fact that it is not straightforward to assess the phenotypic heterogeneity of an isogenic population in a mixed community. In order to circumvent this issue we present a membrane-based synthetic community setup. Using this setup we investigated the effect of microbial interactions on the individual phenotype and phenotypic diversities of the interacting taxa.

### Effect of interaction on phenotype and phenotypic diversity

Based on flow cytometric fingerprinting, the phenotypic diversity of both community members was lower when they were grown in a coculture compared to when they were grown as axenic cultures (**Fig. 2 A, S6 and S7**). This effect of interaction on population diversity was more pronounced for *Enterobacter* than for *Pseudomonas*, indicating that different taxa had different phenotypic responses to the interaction. When comparing the phenotypes of the populations through beta-diversity assessment (**Fig. 3**) and Raman spectroscopy (**Fig. 5**) a similar observation was found. The differences between the phenotypic state of *Pseudomonas* in the coculture and in the axenic culture were smaller compared to the differences between *Enterobacter* in the coculture and in the axenic culture.

Differences in scattering patterns were limited for both taxa, implying that there were no large changes in cell morphology (44). Since SG staining is a stoichiometric staining, a higher fluorescence signal is directly related to a higher concentration of stained nucleic acids (45, 46). In terms of nucleic acid content, large differences were observed for *Enterobacter* and limited differences for *Pseudomonas*, with *Enterobacter* shifting towards high nucleic acid individuals (**Fig. 2 B and C, Fig. S7**). This can indicate different physiological shifts. On the one hand, the DNA copy number could be increased, implying an adaptation of the cell cycle. Although both bacteria were expected to be in stationary phase at all sampling points (**Fig. S1**), it is possible that under stress, the bacteria adapted their cell cycle behaviour and DNA concentration (47). On the other hand, the bacteria might have maintained a similar DNA concentration but a higher RNA concentration, indicating a shift in their gene expression. The bacteria could have been more actively expressing the same genes as they were in the axenic cultures, or they might have shifted towards expression of other genes compared to the axenic cultures. Lastly, also an increased membrane permeability may explain higher fluorescence signals.

Through single-cell Raman spectroscopy, which offers an in depth view on the biochemical phenotype, we attempted to investigate which of the above mentioned scenarios was most likely to be occurring. Using a reference-based peak assignment, the Raman spectra indicated differences in wavenumbers which were potentially related to DNA and RNA, and in this way support both hypotheses (**Fig. S8**). It should be noted that the tentative peak assignment resulted in inconsistent conclusions regarding the intensity change of nucleic acid related wavenumbers for both taxa under the different conditions (*i.e.* axenic or coculture). This might be explained by the fact that the signal intensity at every wavenumber is the superposition of all compounds signals at this wavenumber, thereby prohibiting biomolecule-specific interpretation.

Several uptake- or metabolic pathways are often simultaneously active in a single taxon’s population (48, 49). Interspecies interactions are known to alter the intensity of the production pathways that are active in interacting bacteria (50, 51), and hence, they may be influencing population heterogeneity. For example, the interspecies interactions may allow species to share products of costly pathways, and in this way deprioritize some functions which would be necessary for the proliferation in monoculture, such as production of certain amino acids (50, 52). Since costly production pathways are often expressed by only a fraction of a clonal population (15, 53), sharing of these pathways between genotypes might allow one or both interacting genotypes to steer the distribution of their costly phenotypes, and hence reduce their population heterogeneity. This would enable each genotype to occupy the functions at which it is most performant, thus, creating a mixed community with a higher overall performance. The increased cell density in the mixed culture as compared to the axenic cultures may indicate this increased performance (**Fig. S5**). The idea of pathway sharing is in line with the observation that the gene-essentiality for a specific taxon is dependent on its community partners (54). Asides these cooperative interactions, competition may also explain the reduction in phenotypic diversity. It may confer a competitive advantage for a taxon to reduce its heterogeneity and in that way reduce the fraction of individuals that are in a suboptimal state for exploiting the current environmental conditions (13). In this study, the community was predicted to be dominated by *Pseudomonas* (**Fig. 4**). A possible explanation for the fact that *Enterobacter* showed a stronger reduction in phenotypic diversity may be that *Enterobacter* needed to reduce its heterogeneity more in order to compete with *Pseudomonas*.

### Evaluation of the experimental setup

In literature, phenotypic heterogeneity is most often studied though the assessment of single cell metabolic activity, using isotope labelling with stable or radioactive probes (49, 55), or though the quantification of gene-expression variability with fluorescent labelled proteins (2, 11, 48). Both isotope labelling and fluorescent labelled proteins allow to study heterogeneity in clonal populations. However, they require either a modification of the organisms under study by inserting a fluorescent protein or the use of rather expensive, and sometimes dangerous, isotopes. Using phenotypic fingerprinting through flow cytometry does not require any tagging of bacteria or the use of isotopes. Moreover, it is possible to assess the phenotypic diversity of bacterial populations without prior knowledge on potentially relevant metabolic pathways (isotope labelling) or genes (fluorescent labelling). The main benefits of the flow cytometric approach are its speed and the fact that large amounts of cells can be analysed. This allows to have good coverage of the phenotypic landscape of the community and to achieve a highly resolved sampling frequency.

However, when assessing phenotypic heterogeneity, there needs to be a definition of the phenotypes between which will be distinguished. Using the previously published protocol by Props *et al.*, (2016), a binning grid was applied to each of the bivariate parameter combinations (*i.e.* scatter and fluorescence parameters). Bacteria that fell within the same bin were defined as the same phenotype. Thus, phenotypes, and by extension the phenotypic diversity metrics, were defined *ad hoc.* Moreover, when evaluating phenotypic heterogeneity based on flow cytometry, the phenotypic traits on which information is gained are morphological parameters and nucleic acid content (in case of SG staining). But only a certain level of information is retained in the scatter and fluorescence parameters (*e.g.* morphology cannot be inferred directly from scatter values) (56). Thus the phenotypic traits derived through flow cytometry are an abstract representation of the phenotype. Additionally, only taking into account these traits is an abstraction of the entire phenotypic diversity of the bacteria. The fact that phenotypes were defined using a predefined binning grid and based on a limited number of phenotypic traits, makes it difficult to make a link with functionality and to fully understand the underlying biological or ecological process that caused the phenotypic diversity shift. Additional examination of the transcriptome (52, 57, 58) or exometabolite profiles (59) could provide valuable insights in the cause of the phenotypic adaptation and the functional consequences that the change in phenotypic state might bring. Additionally, more validated and automated pipelines for detection of biomolecules based on single-cell Raman spectra would be an interesting improvement.

### Conclusion

In conclusion, we have used a synthetic community setup in which the individual phenotypic heterogeneity of environmental isolates in mixed or synthetic communities can be studied. We demonstrated that interactions between bacterial populations lead to an adjustment of the individual phenotypic diversities of the interacting populations. As phenotypic heterogeneity is playing an important role in pathogenicity and virulence (14), antibiotics resistance (12, 60), biotechnological applications (20, 23, 61, 62), ecosystem properties (63), it is crucial to understand its influencing factors. The experimental design presented in this study provides a framework within which further ecological hypotheses regarding phenotypic heterogeneity and microbial interactions can be tested.

## Materials and Methods

### Isolates

An *Enterobacter* sp. and a *Pseudomonas* sp. were selected from a set of drinking water isolates which were isolated on R2A agar and provided by Pidpa (Provinciale en Intercommunale Drinkwatermaatschappij der Provincie Antwerpen, Belgium). Preliminary tests showed that these isolates had distinct cytometric fingerprints, as determined by the method of Rubbens et al. (30), and reached stationary growth phase in M9 supplemented with 200 mg/L glucose within 24 hours at 28°C, starting from a cell density of 10^6^ cells mL^-1^ (**Fig. S1**). The isolates were identified with Sanger sequencing (LGC Genomics GmbH, Germany). The strains were deposited into the BCCM/LMG Bacteria Collection under accession IDs LMG 30741 (*Enterobacter* sp.) and LMG 30742 (*Pseudomonas* sp.).

### Experimental setup

Bacteria were cultured in Transwell plates (Corning^®^ Costar^®^ 6-well cell culture plates, Corning Incorporated) where apical and basal compartments were created using cell culture inserts (ThinCert^™^ Cell Culture Inserts with pore diameter 0.4 μm, Greiner Bio-One). The membranes of the culture inserts were replaced by membranes with smaller pore sizes to avoid migration of bacteria between the two compartments (Whatman^®^ Cyclopore^®^ polycarbonate and polyester membranes with 0.2 μm pore size, GE Life Sciences). Four synthetic communities were created, being two axenic cultures, a physically separated culture and a mixed culture (**Fig. 1**). Each community was prepared in triplicate and randomised over the plates to account for plate effects. Before the start of the experiment, both bacteria were grown on nutrient agar (Oxoid, UK) plates. A single colony was picked from each plate and transferred to liquid minimal medium (M9 with 200 mg/L glucose as carbon source). After two days of incubation at 28°C, cell densities in the liquid cultures were determined by flow cytometry and the cultures were diluted to the desired starting cell densities in fresh medium. The required dilution was high enough to neglect differences in volume of fresh medium, and thus resources for growth, that were needed to prepare the cultures. The starting cell densities were set to have the same initial cell density of 10^6^ cells mL^-1^ in each synthetic community, and with equal relative abundances for both community members in the cocultures and mixed cultures (**Table S1**).

The 6-well plates were incubated at 28°C and gently shaken (25 rpm) to aid diffusion of metabolites between the compartments. The communities were monitored over a period of 72 hours. Every 24 hours samples were analysed by flow cytometry. After 72 hours samples were fixed with 4% paraformaldehyde for Raman spectroscopic analysis (Supplementary material and methods). Sample fixation was necessary since single-cell Raman measurements were too time consuming for immediate analysis. The first sampling moment was at 24 h, which suggests, based on the previously determined growth kinetics, that both taxa were in stationary phase at every sampling point (**Fig. S1**).

### Flow cytometry

For flow cytometric analysis, the samples were diluted and stained with 1 vol% SYBR^®^ Green I (SG, 100x concentrate in 0.22 μm-filtered DMSO, Invitrogen) for total cell analysis, and with 1 vol% SYBR^®^ Green I combined with propidium iodide (SGPI, 100x concentrate SYBR^®^ Green I, Invitrogen, and 50x 20 mM propidium iodide, Invitrogen, in 0.22 μm-filtered dimethyl sulfoxide) for live-dead analysis. SG primarily stains double stranded DNA, but will also stain the RNA (31). Staining was performed as described previously, with an incubation period of 20 min at 37°C in the dark (32). Samples were analysed immediately after incubation on a FACSVerse^™^ flow cytometer (BD Biosciences, Belgium), which was equipped with eight fluorescence detectors (527/32 nm, 783/56 nm, 586/42 nm, 700/54 nm, 660/10 nm, 783/56 nm, 528/45 nm and 488/45 nm), two scatter detectors and a blue 20-mW 488-nm laser, a red 40-mW 640-nm laser and a violet 40-mW 405-nm laser. The flow cytometer was operated with FACSFlow^™^ solution (BD Biosciences, Belgium) as sheath fluid. Instrument performance was verified daily using FACSuite^™^ CS&T beads (BD Biosciences, Belgium).

### Raman spectroscopy

Prior to analysis, the fixed sample was centrifuged for 5 minutes at room temperature and 5000 g, and the pellet was resuspended in 0.22 μm-filtered milli-Q (4°C). 10 μL of cell suspension was spotted onto a CaF_2_ slide (Crystran Ltd., UK) and air-dried for a few minutes. The dry sample was analysed using an Alpha 300 R confocal Raman microscope (WITec GmbH, Germany) with a 100x/0.9NA objective (Nikon, Japan), a 785 nm excitation diode laser (Toptica, Germany) and a UHTS 300 spectrometer (WITec GmbH, Germany) with a -60°C cooled iDus 401 BR-DD CCD camera (Andor Technology Ltd., UK). Laser power before the objective was measured daily and was about 150 mW. Spectra were acquired in the range of 110-3375 cm^-1^ with 300 grooves/mm diffraction grating. For each single cell spectrum, the Raman signal was integrated over 40 s. All Raman samples were analysed within 1 week after sampling, with minimal time between them to limit possible differences caused by differences in storage duration. For each population between 51 and 55 single cell spectra were measured from a single biological replicate population. To allow for a fair comparison, 51 spectra were selected from each population for further analysis. The spectra with the lowest intensity were assumed to be of lesser quality, and were therefore discarded.

A large peak in the range of 850 - 1030 cm^-1^ was present in the spectra of *Enterobacter* in the axenic culture, while this peak was not observed in the other populations or during preliminary tests (**Fig. S2**). Moreover, intensity values showed large variability for this region. This might be the result of technical issues during fixation or storage of the sample. Similar to the study of García-Timermans et al. (33), this region was excluded for further analysis (**Fig. S3**).

### Data analysis

#### Flow cytometry

##### Phenotypic diversity analysis

The flow cytometry data was imported in R (v3.3.1) (34) using the flowCore package (v1.40.3) (35). A quality control of the dataset was performed through the flowAI package (v1.6.2) (36). The data was transformed using the arcsine hyperbolic function and the background of the fingerprints was removed by manually creating a gate on the primary fluorescent channels (32). The Phenoflow package (v1.1.1.) (37) was used to assess the phenotypic community structure of the bacterial populations. Based on the study of Rubbens et al. (38), which assessed the usefulness of information captured by additional detectors (*i.e.* detectors that are not directly targeted) for bacterial population identification, an optimal subset of detectors was selected to include in the analysis. The subset included the scatter-detectors, the detector for which had been stained (*i.e.* FITC), and some additional detectors that received spill-over signals (AmCyan, dsRed and eCFP).

Prior to diversity estimation, all populations were subsampled to 30,000 cells in order to account for sample size differences. In short, for each bivariate parameter combination (*i.e.* combination of the scatter and fluorescence parameters) an 128×128 equal space binning grid is applied, which discretizes the parameter space, and in which each bin represents a hypothetical phenotype. For each bin a kernel density estimation is performed. All density estimations are summed to the total density estimation of the community. Finally, the density values for each of the bins are concatenated into a 1D-vector, which is called the ‘phenotypic fingerprint’. From this fingerprint, alpha and beta diversity are calculated, which are used as measures for phenotypic population heterogeneity. The alpha diversity (*i.e.* within sample diversity) is calculated by means of the first three Hill diversity numbers D_0_, D_1_ and D_2_, which correspond to the observed richness, the exponential of Shannon entropy, and the inverse Simpson index, respectively (39). Beta diversity (*i.e.* between sample diversity) is evaluated by principal coordinate analysis (PCoA) on the Bray-Curtis dissimilarities between the fingerprints. Significance of the differences between fingerprints was assessed by means of PERMANOVA on the Bray–Curtis dissimilarity matrix. Homogeneity of variance in groups was assessed before performing PERMANOVA. A significance level of 0.01 was used.

##### In silico communities

Relative abundances in the mixed cultures were predicted using the supervised in silico community methodology of Rubbens, Props, Boon, *et al.* (2017), implemented in the Phenoflow (v1.1.1) software package. In short, a cytometric fingerprint of the taxa that make up the synthetic community is made. Next, the single-cell data of the axenic cultures is aggregated to a so-called ‘in silico community’. This in silico community consists of labelled data, which allows the use of supervised machine learning techniques, such as random forests, to discriminate between different community members. The label to be predicted is the taxon and the predictors are the scatter and fluorescence parameters. Once this classifier has been trained on the dataset, it can use the single-cell data to predict the relative abundances of both taxa in a mixture. For training of the random forests, the biological replicates were pooled together and 10,000 cells of both *Enterobacter* and *Pseudomonas* were randomly sampled.

##### Raman spectra

The data was analysed in R (v3.3.1). Spectral processing was adapted from the study of Berry et al. (40), and was performed using the MALDIquant package (v1.16) (41). In short, baseline correction was performed using the statistics-sensitive nonlinear iterative peak-clipping (SNIP) algorithm. Next, the biologically relevant part of the spectrum (600-1800 cm^-1^) was selected (25) and the spectra were normalised by surface normalisation. The intensity values were zero centred and scaled to unit variance before performing PCA (stats package, v3.3.4).

### Data availability

The entire data-analysis pipeline is available as an R Markdown document at https://github.com/jeheyse/Cocultures2018. The Raman data and accompanying metadata are available at https://github.com/jeheyse/Cocultures2018. Raw FCM data and metadata are available on FlowRepository under accession ID FR-FCM-ZYWN (for review: https://flowrepository.org/id/RvFrlZ3CQpF6XTlkKEtSHYE9VPTRoJREiYJJz8HKfdO9nITuTMc2JA3HiXvPt5fE).

## Acknowledgements

The authors want to thank Katrien De Maeyer (Pidpa) for providing the isolates. We thank Tom Bellon for his assistance during the flow cytometric measurements and Dmitry Khalenkow for his assistance in the Raman spectroscopy measurements. This work was supported through the Geconcentreerde Onderzoeksactie (GOA) from Ghent University (BOF15/GOA/006). JH is supported by the Flemish Fund for Scientific research (FWO-Vlaanderen, project 1S80618N). BB is supported by the Institute for Innovation by Science and Technology in Flanders (IWT, project 131370). RP is supported by Ghent University (BOFDOC2015000601). PR is supported by Ghent University (BOFSTA2015000501).

## Contributions

JH, RP, PR, BB, AS, WW and NB conceived the study. JH carried out the laboratory work and analysed the data. JH, BB, RP and PR interpreted the results and wrote the paper. NB, WW and AS supervised the findings of this work. All authors reviewed and approved the manuscript.

